# Bird community structure in a mixed forest-production landscape in the northern Western Ghats, India

**DOI:** 10.1101/2022.04.04.486917

**Authors:** Avishkar Munje, Ajith Kumar

## Abstract

Production landscapes outside protected areas are important for the conservation of wildlife, especially in countries like India with high biodiversity and human densities. Production landscapes like plantations often occur in close proximity to biodiversity-rich areas. Rubber and cashew are lucrative plantation crops in India, which although grown under similar environmental conditions, vary in their management and productivity. These plantation crops are often found along the edges of forests, thereby creating a buffer between forested and urban landscapes. While these areas have the potential to be a refuge for species otherwise restricted to natural forests, they are poorly-studied. To address this knowledge gap, we studied how habitat type (cashew, rubber or forest) and habitat characteristics affect bird diversity and guild structure in the Tillari landscape of Western Ghats, Maharashtra. Additionally, we examined how these effects are mediated by distance of plantations to nearest forest edge. In each habitat type, in 2018, we sampled birds six times each in 30 locations using fixed radius point counts. We found that bird diversity in cashew plantations (14 species) was comparable to that of adjoining forests (15 species). Rubber plantations, on the other hand, had lower bird diversity (9 species) than that in cashew or forests. When bird diversity was analysed based on dietary guilds, rubber plantations had fewer bird species in all guilds and cashew plantations had higher abundance of nectivores and lower richness of insectivores than in forest. Distance from the forest did not affect bird diversity in rubber plantations, whereas cashew plantations had fewer nectivorous birds and higher insectivorous birds away from the forest edge. Our results show that cashew plantations can serve as an important surrogate habitat for forest birds in the northern Western Ghats. The findings indicate the unsuitability of rubber plantations for sustaining bird diversity. While there are many studies available on bird diversity in rubber plantations, this is among the first studies of bird community structure in cashew plantations. At a time when forests are rapidly being cleared for plantations, our findings provide valuable data to examine the differential impacts of plantation type on biodiversity.

## INTRODUCTION

Terrestrial protected areas (PAs) occupied only about 13% of global land area as of 2011 (Convention on Biological Diversity, 2011). The last two decades have seen very little improvement in PA coverage of threatened mammals, birds and amphibians, with nearly 17% of these species still occurring outside protected areas and 85% of them inadequately covered within existing protected areas (Venter et al., 2014). The Aichi 2020 targets under the Convention of Biological Diversity are difficult to achieve due to the huge lost-opportunity cost to agriculture. Further, the proposed increase in PA coverage would only result in a marginal increase in the coverage of threatened species (Venter et al., 2014). Since the scope for increasing PA coverage is very limited (Jenkins and Joppa, 2009), there is an urgent need to consider other forms of land use as conservation opportunities.

A recent study shows that nearly 27% of the global forest loss between 2000 and 2015 was permanent and due to commodity production, including plantation crops such as rubber and oil palm (Curtis et al., 2018). Other forms of forest loss such as shifting cultivation, forestry and wildfire are temporary in nature (Curtis et al., 2018). Production landscapes cover 38% (Foley et al., 2011) of the earth’s landmass and are known to support a wide variety of taxa, housing ecologically sensitive species in the tropics (Bhagwat et al., 2008 for a review; Karanth et al., 2016). The survival of many species would depend on their ability to survive in different land-uses (Benton et al., 2003; Bhagwat et al., 2008; Sodhi et al., 2010; Phalan et al., 2013).

Studies have shown that commodity plantations can vary considerably in the biodiversity that they harbour. Oil palm supports very little biodiversity (Koh and Wilcove, 2008). Rubber plantations also support only a fraction of species found in neighbouring forested landscapes (Li et al., 2007) and this is also true for tea plantations (Kottawa-Arachchi and Gamage, 2015). In contrast, coffee plantations can harbour a substantial component of biodiversity found in adjoining forests (Bhagwat et al., 2008).

In general, the presence of remnant forest patches is a major determinant for the occurrence of biodiversity in human-modified landscapes in the Western Ghats (Anand et al., 2010). A spill-over effect is usually seen, with higher species diversity closest to patch edges (Blitzer et al., 2012). Thus, landscapes with both forests and agroforestry plantations could potentially be managed to sustain biodiversity (Chazdon et al., 2009). Human-use landscapes that are well-managed in favour of biodiversity could serve as high quality habitats and matrices that can facilitate biodiversity dispersal across fragmented landscapes (Perfecto and Vandermeer, 2008). There is thus a growing need to view forested areas and human-use areas as interacting entities of a single system, rather than as independent components (Gardner et al., 2009).

Past studies in cashew have looked at mammalian diversity (Rege 2016) human primate interactions (Hockings et al., 2012, 2013; Casanova et al., 2014), butterfly diversity (Vasconcelos et al., 2015; Mahata et al., 2019) and spider diversity (Bhat et al., 2013). However, there have been no studies on birds so far.

The Western Ghats in India, with Sri Lanka, is one of the 25 global biodiversity hotspots (Myers et al., 2000). This area also has high local endemism, making it a high priority region for conservation and establishment of protected areas (Bossuyt et al., 2004; Bhagwat et al., 2005). Of the 160,000 km^2^ expanse of the Western Ghats, natural habitats cover about one-third of the land, with the PA network spanning 58 parks and 13,595 km^2^ (Anand et al., 2010). India perhaps has very limited scope in increasing PA coverage, which has remained at about 5% (compared to 13% globally) for the last two decades (Ghosh-Harihar et al., 2019). It is a landscape where natural, semi-natural and agroforestry systems occur in close proximity forming a unique mosaic of interacting components (Anand et al., 2010). Two-thirds of land is spread across diverse uses including reservoirs, open agriculture, human habitations and plantations of coffee, tea, rubber and cardamom, interspersed within other cash crops (Daniels et al., 1990). These commodity plantations span over 10,000 km^2^ of the Western Ghats (Anand et al., 2010). Tea plantations occupy another 1,100 km^2^ of the high-rainfall regions and require little to no shade, thereby standing out in stark contrast to its surrounding forests structurally and ecologically (Tea Board of India, 2009). Rubber, sometimes intercropped with pineapple (*Ananus comosus*), banana (*Musa spp*.), turmeric (*Curcuma longa*), ginger (*Zingiber officinale*) and others, span 5000 km^2^ in area (Rubber Board of India, 2003).

Coffee plantations, which are almost entirely shade-grown in the Western Ghats, occupy over 3000 km^2^ (Coffee Board of India, 2009). Coffee is grown under shade of native or non-native tree species like silver oak. Large proportions of non-native trees have been shown to negatively affect overall bird diversity in coffee plantations. Plantations further away from forests show a decline in range-restricted species of birds (Anand et al., 2008), and an overall decline in mammalian diversity (Bali et al., 2007) and butterfly diversity (Dolia et al., 2008). These plantations can also act as a buffer and facilitate movement of animals between different forests (Chazdon et al., 2009). While coffee supports substantially higher bird diversity than areca and rubber (Karanth et al., 2016), all these agroforests play an important role in providing additional refuges for wildlife in the Western Ghats.

Much of the Western Ghats are unprotected or private forests, where rubber and cashew are important cash crops to plantation farmers. As of 2017-18, 4700 km^2^ was under cashew plantation with a production output of 4,80,000 metric tonnes (Directorate of Cashew and Cocoa Development, states of Maharashtra, Kerala, Goa, Karnataka). Cashew was introduced to India in the 16^th^ century, but its economic importance was realized in the early 20^th^ century. The first rubber plantation in this region was in the 1970s. Cashew as a crop requires lower maintenance (and therefore less human intervention) than rubber. While fruit harvest for cashew takes place during only two months of the year, rubber is tapped year-round. Cashew plantations in the landscape are either privately-owned or community-owned. Based on the nature of ownership and topography of the land, the plantations have a varying degree of vegetation structure and composition. While community plantations are seen only along the hillside (mostly bordering private forests), private plantations are found on slopes as well as plains. Cashew plantations on slopes, both private and community owned, have some amount of native vegetation as trees and understory (weed and shrubs), but plantations away from the hills have few native trees and no understory.

Rubber plantations are a more recent phenomenon. These plantations vary from a few hectares to more than 1000 hectares in area, with some of large plantations adjoining reserve forests, occurring on both hilly and flat terrain, and have been created on farm land or private forests. Rubber is characterized by a uniform girth and height class of tree, no understory except for one leguminous creeper (*Mucuna bracteata*), and interspersed occasionally with small patches of banana and areca. These plantations are intensively managed and tapped for rubber and some are electric fenced.

Despite cashew and rubber plantations being a major land use in the Western Ghats, often adjoining natural forests, their potential for conservation of biodiversity has been very little studied. Rege (2016) found that of 11 large mammals found in the forests of northern Western Ghats, nine were also found in adjoining cashew plantations. We were thus interested in comparing bird diversity, i.e. species richness and abundance, in cashew and rubber plantations with that of forests in the same landscape.

Bird communities are an important and species-rich taxon in tropical forests. Birds maintain crucial ecological processes like seed dispersal, which ultimately contribute to forest health, and are thus a good focal taxon for conservation assessment and monitoring (Daniels, 1989). In production forests like coffee and cashew the presence of tree cover provides habitat for birds and are therefore better systems of land use for bird diversity than intensive agriculture or pastures. The complexity of habitat (through vertical stratification) and the composition of habitat (through vegetation types and insect abundance) can have predictable links to bird species assemblages with close habitat and dietary associations, especially in species-rich tropical forests (Furness and Greenwood, 1993). Vagile taxa like birds can persist without breeding in production landscapes, but such persistence is likely a function of landscape-level factors like proportion and/or distance of forest in the vicinity. For these reasons, birds are an ideal taxon to study biodiversity in production landscapes (Raman and Sukumar, 2002).

Given the contrasting vegetation structure and management of cashew (*Anacardium occidentale*) and rubber (*Hevea brasiliensis*) plantations compared to forest in this area,we asked -1) How do bird species richness and abundance in cashew and rubber plantations compare with that in forests in the same landscape? 2) How do vegetation attributes influence bird species richness and abundance in these plantations? 3) How do these patterns vary with distance to the nearest forest edge in rubber and cashew plantations?

Given the structural simplicity of rubber compared to cashew, we expected bird diversity to be lower in cashew compared to forest but significantly lower in rubber compared to forest. We expected bird diversity to be lower in habitats with shorter and smaller trees with similar patterns of decline in major dietary guilds and with greater decrease in insectivore and frugivore abundance in rubber (Aratrakorn et al., 2006). We expected a decline in diversity with distance from the nearest forest edge for both plantation crops.

## METHODS

### Study site

I conducted this study in the Tillari landscape of Dodamarg *tehsil*, Sindhudurg District of Maharashtra which forms the southernmost part of what is considered the northern Western Ghats (15°37ʹ to 15°60ʹ north and 73°19ʹ to 73°40ʹ east). Elevation here varies from 50 m to over 1000 m. The average annual rainfall here from the southwest monsoon winds is about 3500 mm. Annual temperature ranges from 12°C to 40°C.

The original vegetation is primarily tropical moist deciduous in the lower elevations and semi-evergreen in the higher elevations (Champion and Seth, 1968). Tree community in the moist deciduous forests consisted of *Tectona grandis, Terminalia bellirica, Syzygium cumini, Garcinia gummi-gutta, Schleichera oleosa* and *Hydnocarpus pentandra* and riparian species such as *Homonoia riparia* and the liana *Entada rheedii*. Recently, a small patch of *Myristica* swamp has also been discovered in the area, which happens to be the northernmost record of this unique habitat type (Sreedharan & Indulkar, 2018). Indian gaur, barking deer, sambar, wild pig, wild dog, leopard and two species of otters along with tigers and elephants are the large mammals reported from the landscape. The region and its surroundings is the northernmost limit for several Western Ghats endemic bird species like Wynaad Laughingthrush, Flame-throated Bulbul, Southern Hill Myna, Grey-headed Bulbul, White-bellied Treepie, Ceylon Bay Owl and Legge’s Hawk-Eagle (Rasmussen and Anderton, 2005). There were about 60 villages in the Dodamarg *tehsil* and human population density according to the latest census was 98 per km^2^ (“Dodamarg taluka”, 2011). Cereal crops (largely paddy cultivation) and plantation crops such as cashew, rubber, pineapple, banana, coconut and areca are the major land-use types in the landscape. Villagers also depend on the forest for cattle grazing, firewood and other forest produce.

Cashew, introduced to the region in the late 16^th^ century by the Portuguese, has been commercially grown in the region only since the early 20^th^ century. Rubber plantations started in the late 20^th^ century here. Rubber plantations are privately owned and range in area from <1 ha to more than 600 ha, are on the slopes and in the plains and are often electric fenced, intensely managed and devoid of any natural vegetation or understory (except for *Mucuna bracteata*, a leguminous creeper planted as a nitrogen fixer). Cashew plantations are grown on community land (mostly on slopes) or private land (in the plains). There is a gradient in natural vegetation in the cashew plantations varying from almost non-existent in intensely managed plantations (mostly in the plains) to being interspersed with private forests on the slopes. Cashew is harvested once a year in March-April while rubber is tapped infrequently throughout the year.

I estimated bird species richness and abundance using point count method, with 30 points each in forest, cashew and rubber (Bibby et al., 2000). These points were sampled 6 times over a period of five months. We estimated pooled species richness and average abundance at point, which were the response variables. We also measured several attributes of vegetation and distance to the nearest forest edge for each point as co-variates. Distance of sampling points in cashew and rubber were obtained using an Etrex Garmin 30x GPS unit.

## Methods

I estimated bird species richness and abundance in the three land uses from point counts which were repeated six times over the study period from January to May 2018. We also estimated habitat attributes from these point count locations, and distance to the nearest forest from digital data.

### Selection of point count sites

I conducted point counts of birds from 30 points each in forest, cashew and rubber. Effort was made to choose points across the entire area, in order to be representative of the landscape and the points were marked using a handheld GPS device (Garmin Etrex 30). Points were chosen on field by taking the vegetation type, aspect of land and percentage of natural vegetation into consideration and marked out using a GPS device. A distance of 250 m was maintained between two points to minimize double counting and all points were 30-50 m from the boundary of the habitat to lower the chances of edge effects.

### Bird sampling

Bird sampling was conducted between January 2018 and May 2018. Birds were sampled using fixed radius point-count method (Bibby et al., 2000). Olympus 10×50m binoculars were used to observe birds and their distance from the sampling point was estimated using a laser rangefinder. After a wait duration of 2 minutes to allow birds to settle down, species identity and abundance of all birds, either seen or heard were recorded within a 7-minute window (Karanth et al., 2016). While the distance of each bird to the centre of the point count was estimated, all detections were restricted to a 30 m fixed radius and then pooled. Birds which could not be identified to species due to poor visibility were identified to genus or family level (less than 1% of the total encounters). Each point was visited six times (three times each in the morning and evening) during the study period, with an interval of at least 15 days.

### Vegetation sampling

At each point-count location, three 5 m x 5 m vegetation plots were placed 5 m from the point location and 120° apart. In each plot, all trees >10cm girth were recorded, its girth at breast height (GBH) using a tape measure and height using a laser range finder were measured. For each point count location in cashew and rubber, distance to the nearest forest edge was calculated on QGIS 3.4.1., using a forest layer created by Mr. Girish Punjabi an ecologist working in Tilari, in 2017, which was manually digitised on ground using a handheld GPS unit.

### Data Analysis

In order to assess sampling adequacy, we made individual-based rarefaction curves for the three habitat types using R Studio. Individual based rarefaction was chosen since the number of sample points was consistent across the three habitats.

For analyses, we used encounters within a 30 m fixed radius as a measure of abundance because detection curves for estimating densities could not be constructed. Additionally, detections were relatively high across all distance bins within the fixed radius. All explanatory variables were standardised (mean = 0, standard deviation = 1) to allow comparison of model parameter estimates. Forest was used as intercept in the model analysis.

Bird abundance for each point was estimated as the average number of birds encountered across 6 temporal replicates. Pooled species richness for each point was calculated as the total number of bird species at a point. We followed Ali and Ripley (1978) for assigning guilds to each species recorded (Ali and Ripley, 1978).

In addition to overall pooled species richness and abundances, we also estimated these parameters for frugivores, insectivores and nectivores in the same manner, using Ali and Ripley (1978) for assigning species to one of these guilds. While encounters of insectivores, frugivores and nectivores were sufficient across habitat types to allow for comparisons, encounters of carnivores, granivores and omnivores were insufficient, and therefore excluded from analysis at guild level.

In order to understand what factors influenced the total and guild-based abundance and richness of birds, we modelled these variables as a function of habitat type (rubber, cashew, forest) and the vegetation attributes that we measured. We used Pearson’s correlation coefficient to test for collinearity among explanatory variables (Fig. S1). If the correlation coefficient between a pair of covariates was greater than |0.4|, only one (which we deemed to be ecologically more relevant) was included in the model. In this case, only tree height and tree girth were selected, while distance to forest edge was modelled separately. We used a generalized linear model (GLM) framework with a Poisson error to model each of the 8 response variables - total abundance, total species richness, guild-based abundance (3 guilds), guild-based species richness (3 guilds) with a. Habitat; b. habitat and tree height; c. Habitat and tree girth; d. Habitat, tree height and tree girth (32 models). To test the effect of distance to the nearest forest edge, we modelled bird abundance and richness for cashew and rubber against distance to the nearest forest edge (4 models). In all the models, forest was modelled as the intercept. Statistically significant trends from each of the models were summarized.

## RESULTS

A total of 14 rubber plantations, 22 privately-owned cashew plantations and one community-owned cashew plantation were sampled in study.

### Bird species richness and abundance across habitat types

In total, 2,837 birds belonging to 99 species and 41 families were recorded, with highest richness observed in forests (76), followed by cashew (62) and rubber (58). Individual-based rarefaction curves indicated sampling completeness in cashew and possible sampling inadequacy in rubber and forest (Fig. 2).

**Fig. 1.**
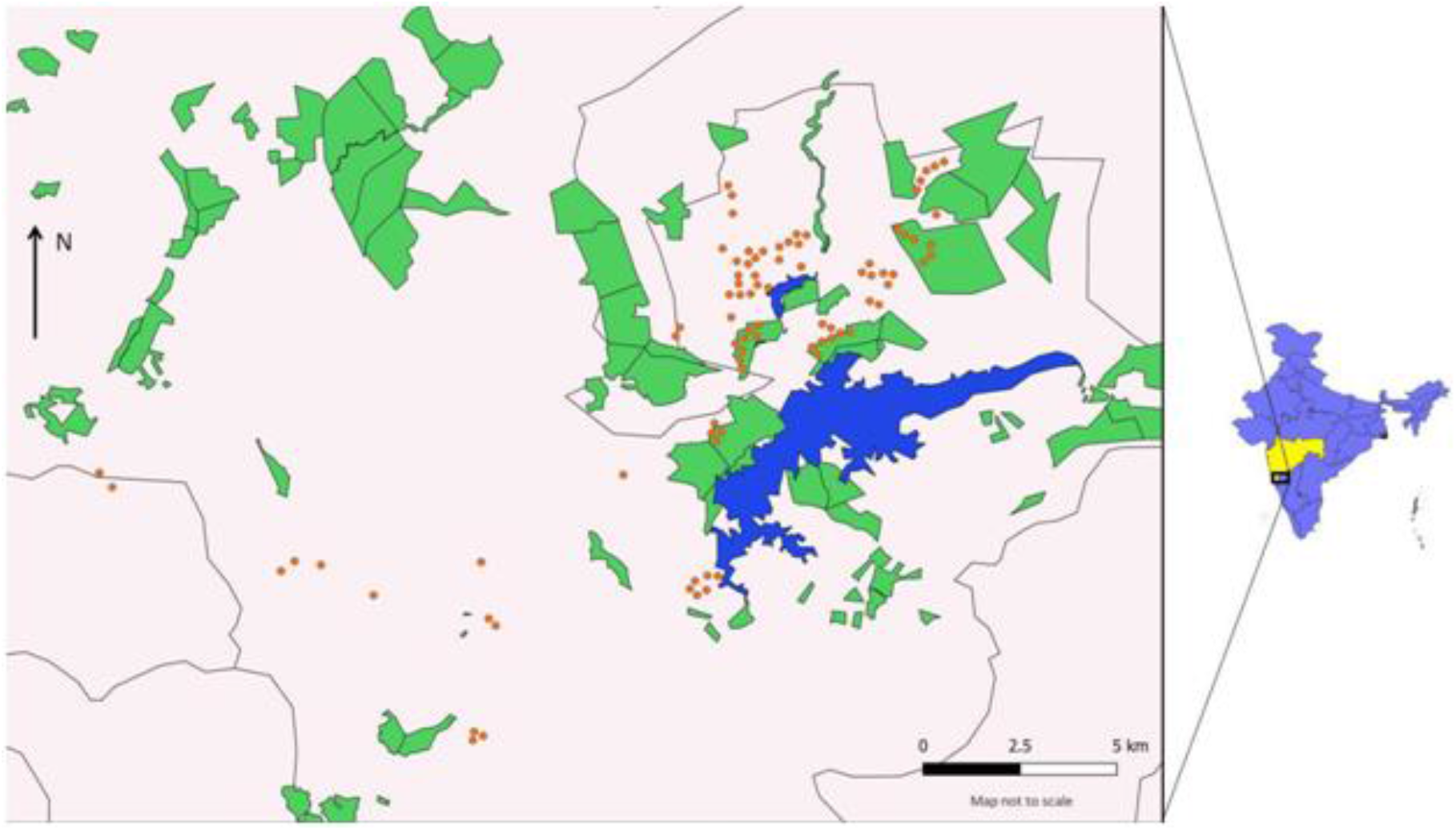
Map of the study area. Sampling locations are represented by orange dots; Reservoir and Reserve forests are represented in blue and green respectively.

**Fig. 2.**
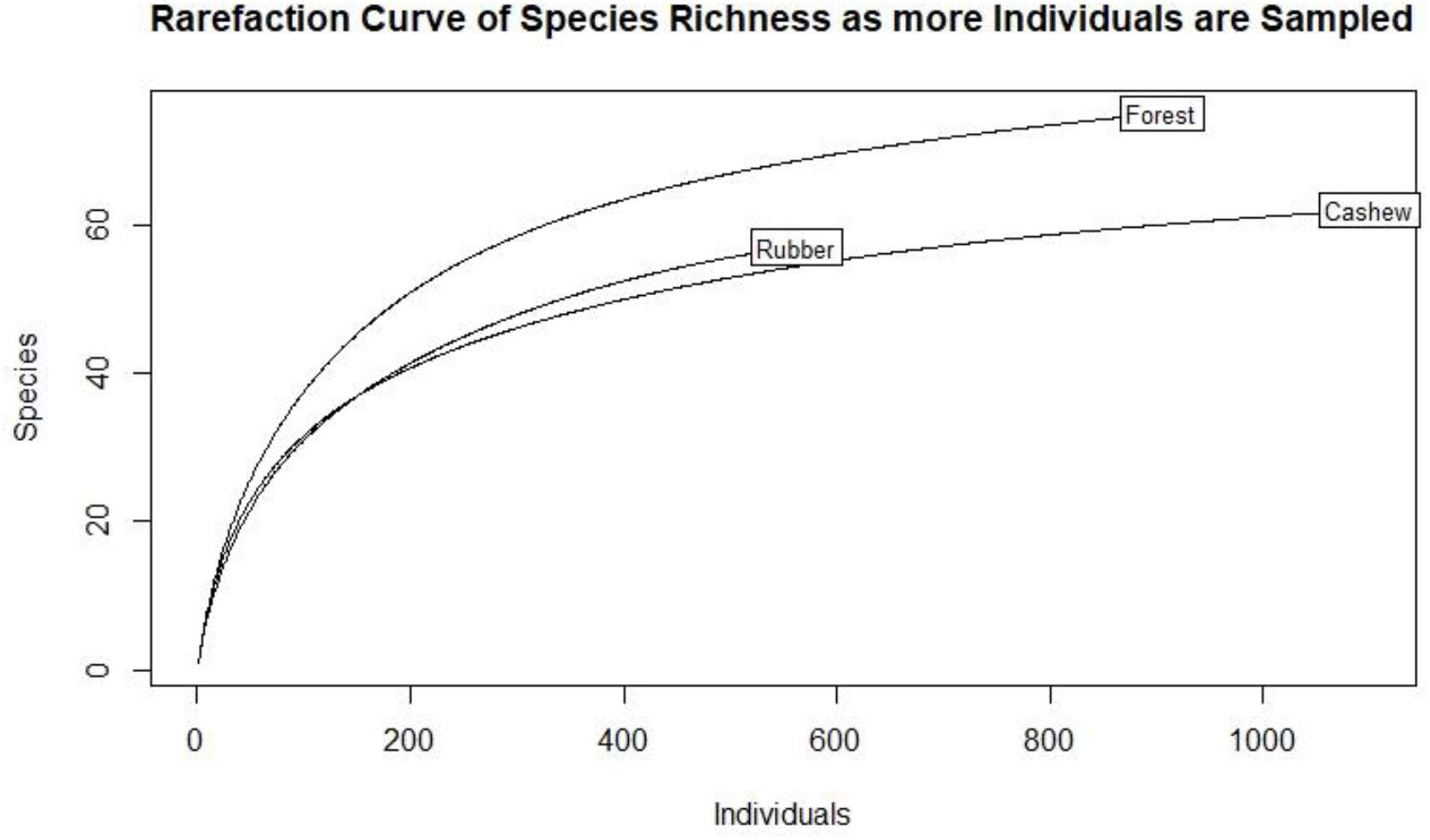
Individual based rarefaction curves for species richness across forest, cashew and rubber.

Cashew and rubber showed similar rarefied species richness up to 200 individuals, beyond which rubber showed greater richness. Forests on the other hand, consistently exhibited higher rarefied species richness than cashew and rubber.

Out of 99 recorded species, 22 species were found only in forests, 9 species only in cashew and 8 species only in rubber. Interestingly, out of 12 recorded Western Ghats endemics, 10 species were found in forest, 7 species in rubber and 2 species in cashew (Table S2). Thirty points were sampled per habitat type and each point visited six times. Mean number of individuals per point was highest in cashew (6.69 ± 0.46) followed by forests (5.71 ± 0.31) and rubber (3.36 ± 0.22). However, forests showed highest pooled species richness per point (15.1 ± 0.68) followed by cashew (14 ± 0.76) and rubber (9.03 ± 0.53) (Figs. 3 and 4).

**Fig. 3.**
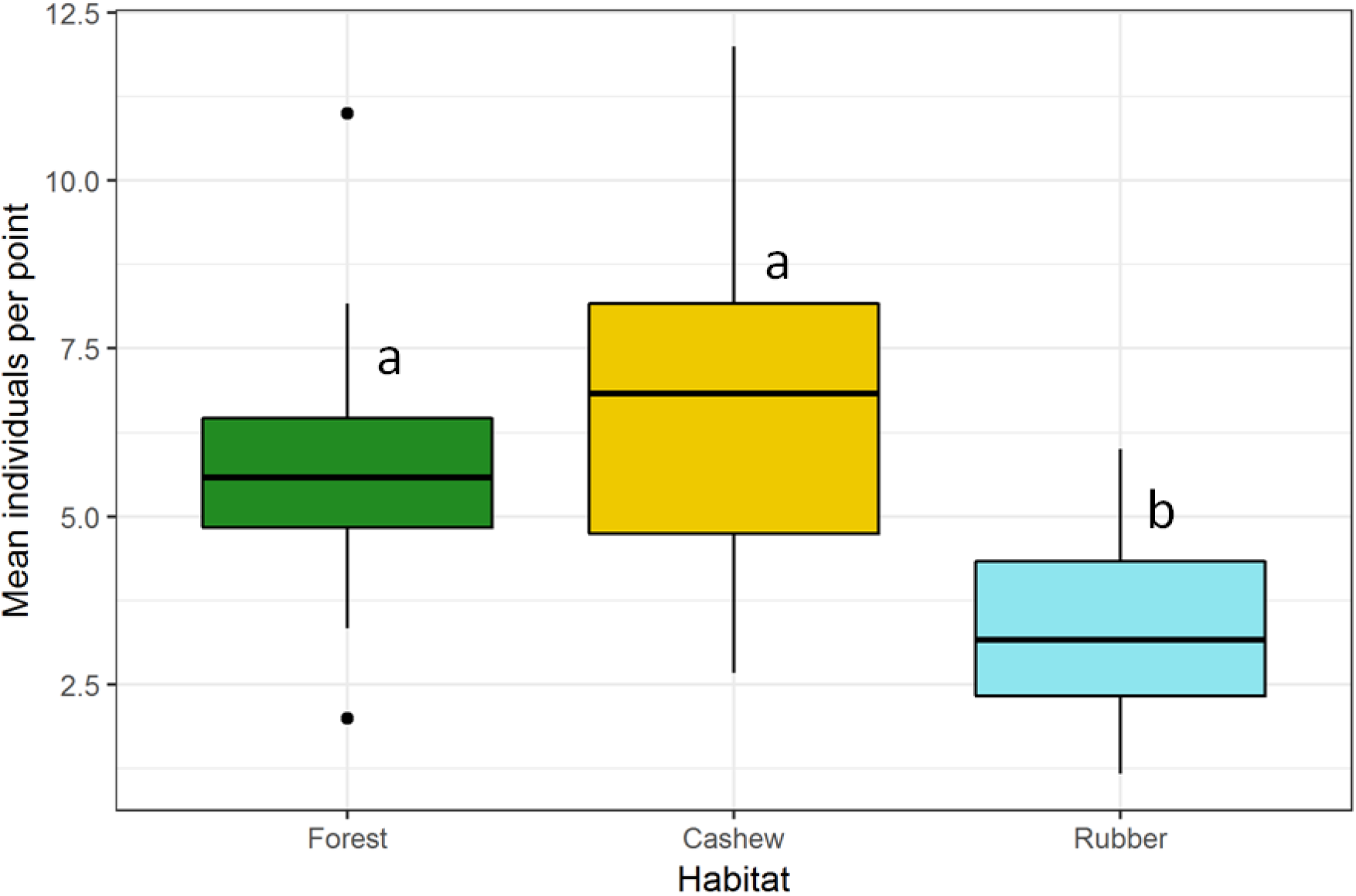
Mean number of individuals per point in the three habitat types. The box covers the inter-quartile region, where the central line inside the box represents the median and the whiskers indicate the remaining 25% of the spread. Dots outside the whiskers are outliers. Same alphabets above the box plot indicate no statistically significant differences (Tuckey HSD test on ANOVA of groups).

**Fig. 4.**
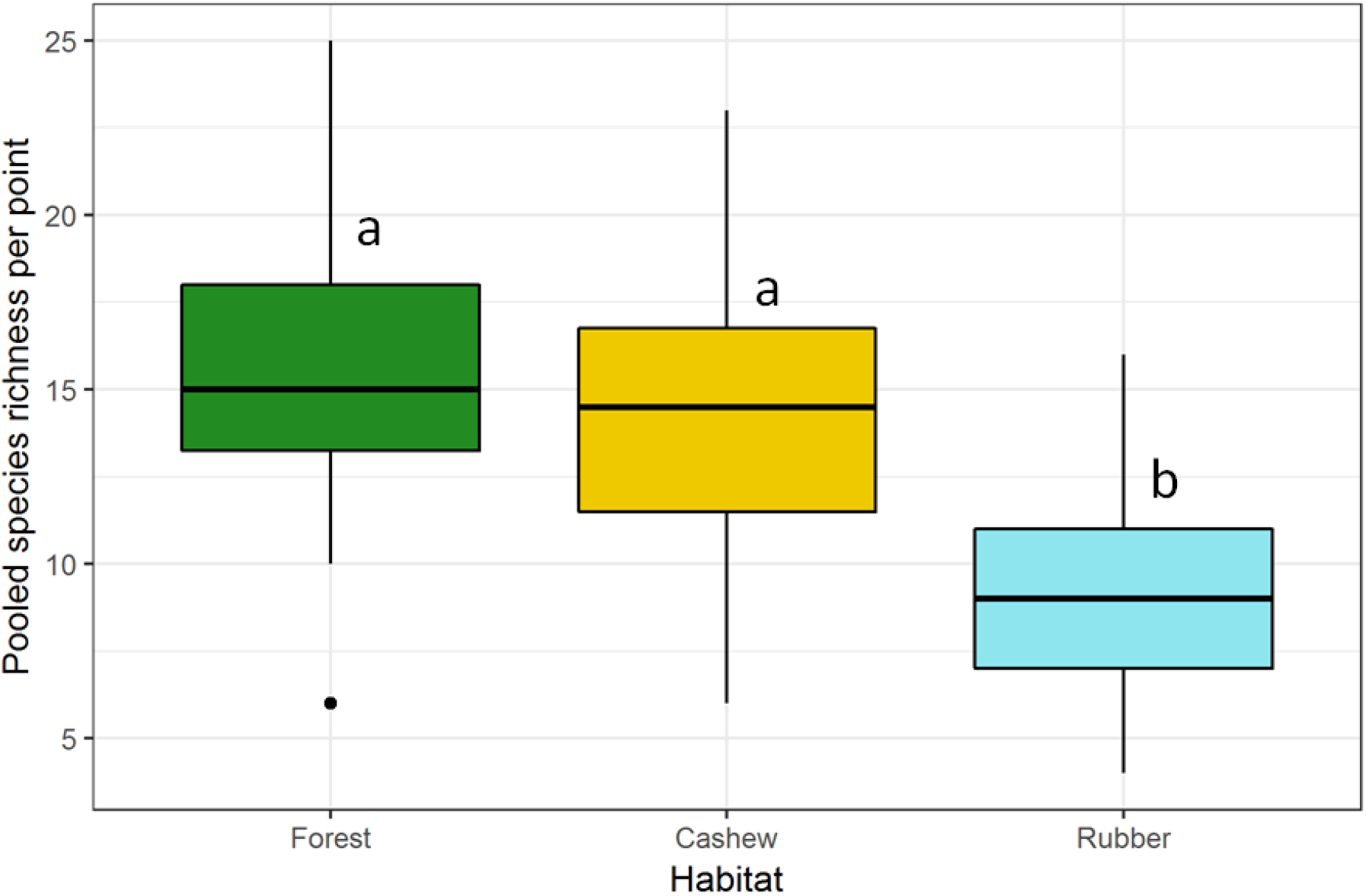
Pooled species richness per point in the three habitat types. The box covers the inter-quartile region, where the central line inside the box represents the median and the whiskers indicate the remaining 25% of the spread. Dots outside the whiskers are outliers. Same alphabets above the box plot indicate no statistically significant differences (Tuckey HSD test on ANOVA of groups).

### Guild structure across three habitats types

Frugivores and nectivores were more abundant in cashew plantations (2.67 ± 0.15 and 2.42 ± 0.2 mean number of individuals per point)) as compared to forest (2.56 ± 0.13 and 1.83 ± 0.09) and rubber (2.24 ± 0.12 and 1.29 ± 0.08). While, nectivores showed the same trend for species richness i.e. highest species richness per point in cashew followed by forest and rubber, frugivores showed highest species richness in forests (5.17 ± 0.27) compared to cashew (4.27 ± 0.27) and rubber (3.50 ±0.22). Insectivores, showed both highest abundance and species richness in forests (3.1 ± 0.19 and 7.50 ±0.52) followed by cashew (3.11 ± 0.28 and 6.33 ± 0.54) and rubber (1.95 ±0.39 and 3.48 ± 0.28) (Figs. 5 and 6).

**Fig. 5.**
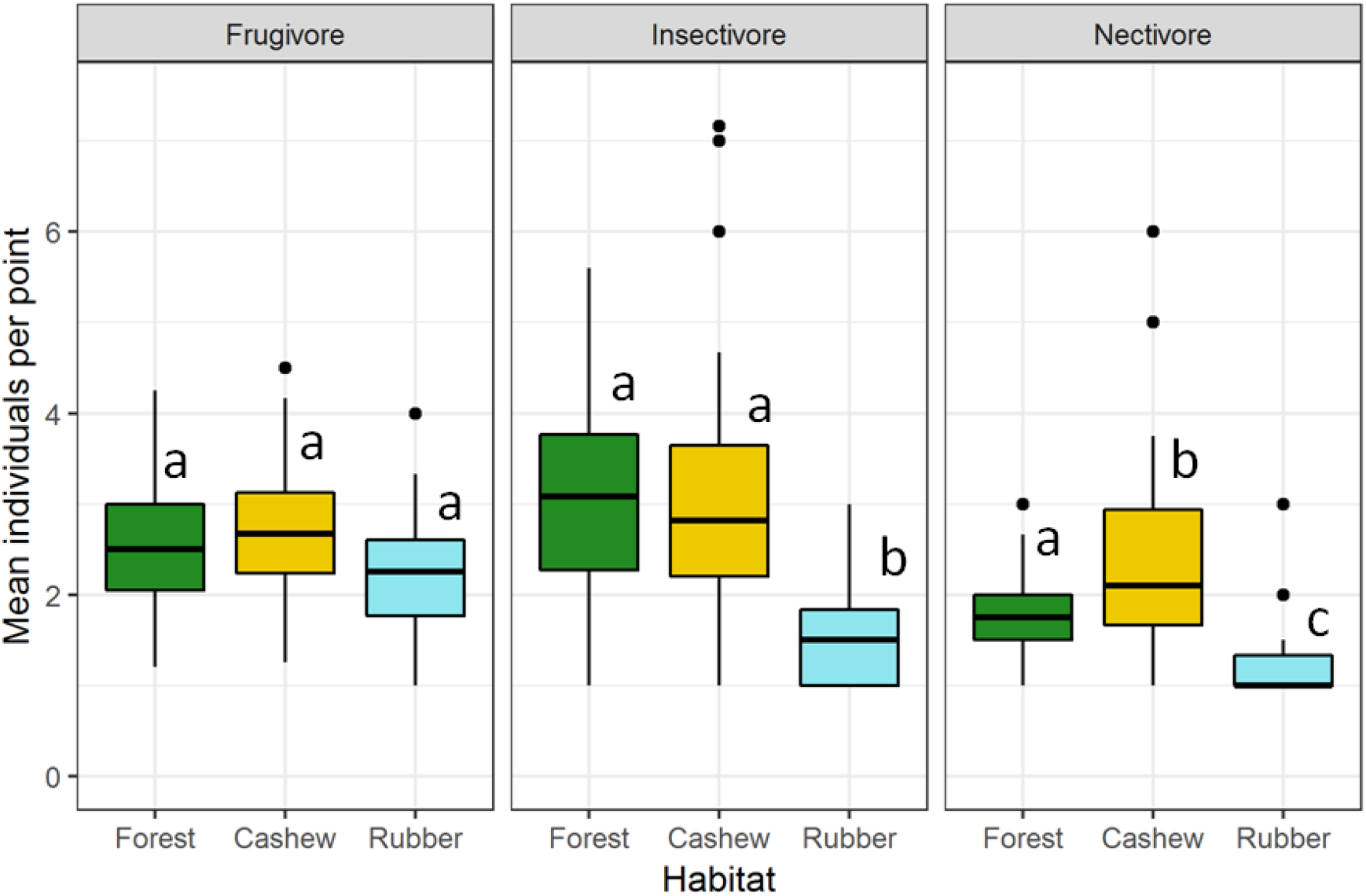
Mean number of individuals of each guild per point for each habitat type. The box covers the inter-quartile region, where the central line inside the box represents the median and the whiskers indicate the remaining 25% of the spread. Dots outside the whiskers are outliers. Same alphabets above the box plot indicate no statistically significant differences (Tuckey HSD test on ANOVA of groups).

**Fig. 6.**
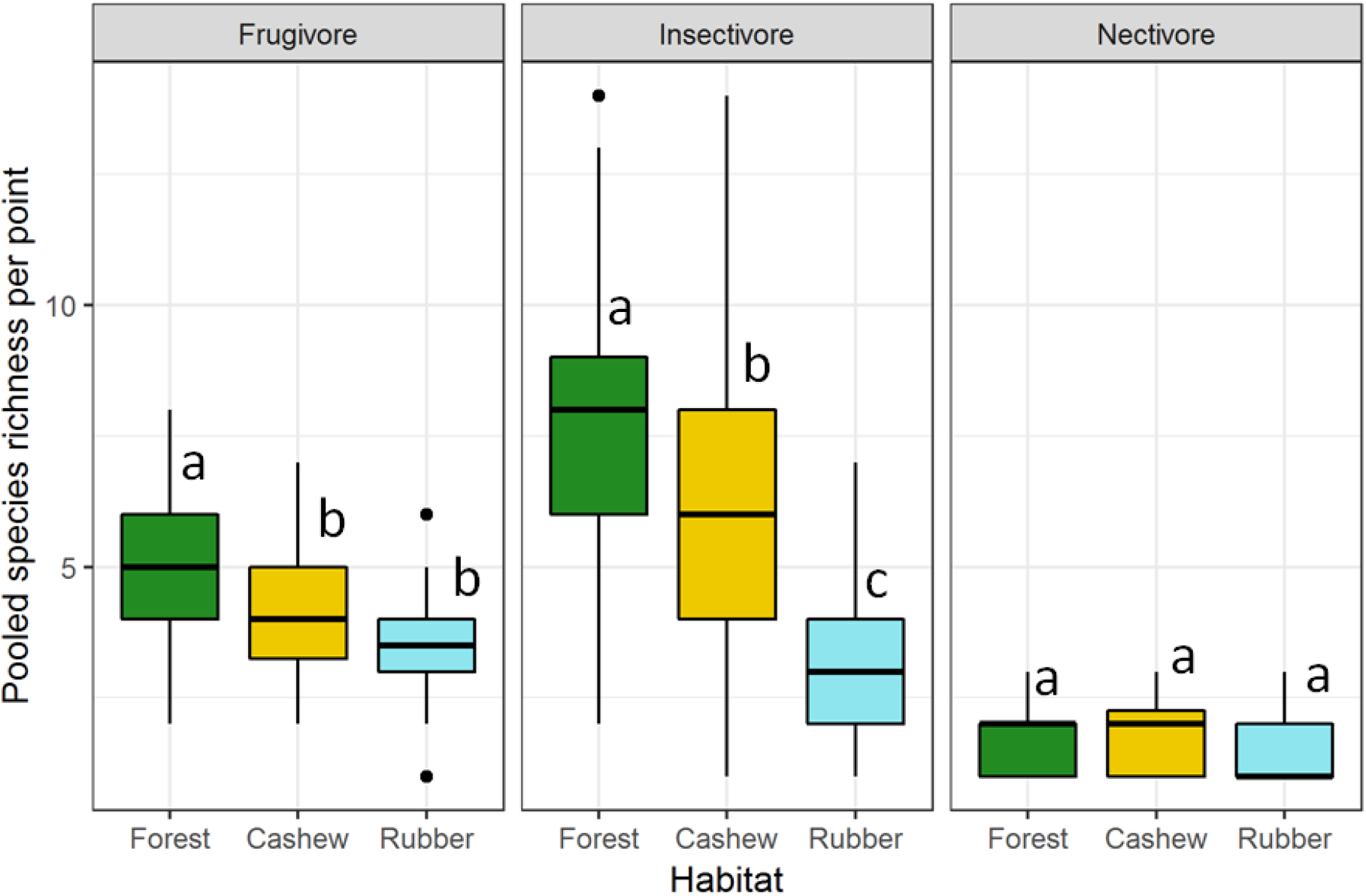
Pooled species richness of each guild per point for each habitat type. The box covers the inter-quartile region, where the central line inside the box represents the median and the whiskers indicate the remaining 25% of the spread. Dots outside the whiskers are outliers. Same alphabets above the box plot indicate no statistically significant differences (Tuckey HSD test on ANOVA of groups).

### Influence of habitat covariates on richness and abundance

#### Abundance parameters

All four models for total abundance (Table 3), showed that total abundance in rubber is significantly lower than that in forest (p<0.001). Two models, one with habitat and tree girth as covariates, and the other with habitat, tree girth and tree height as covariates, showed that total abundance in cashew is significantly higher in cashew as compared to that in forest (p<0.05). Abundance of insectivores was markedly lower in rubber than that in forest (p<0.01). Likewise, abundance of nectivores was lower in rubber (p<0.05) and higher in cashew as compared to that in forest (p<0.05). Tree girth and tree height did not have significant influence on any of the response variables. However, distance to the nearest forest edge showed a positive effect on insectivore abundance in cashew (p<0.05) and a negative effect on nectivore abundance in cashew (p<0.05) (Table 4).

**Table 1.** Predictor variables to be included in the models and the rationale for their inclusion. Of these, only tree height, tree girth and distance to the nearest forest edge were used.

| Covariate | Description |
| --- | --- |
| Canopy cover | Canopy cover affects the amount of light reaching the forest floor which affects the composition and structure of the lower vegetation layers and the species that reside in them. |
| Distance to the nearest forest edge | Closer a plantation is to the edge of the forest, greater is the possibility of finding a number of forest species. |
| Tree density | Plots with dense vegetation could provide good protection and resources for various species |
| Tree girth | Higher is the proportion of larger trees, more is the number of large roosting birds it can support. Bigger trees also tend to harbour more insects for insectivorous species. |
| Tree height | Important for canopy dwelling species. Taller forests have more vertical strata that and could support many niches. |
| Understory | Besides supporting various shy or skulking species, the understory also harbours a variety of resources which includes insects and flowering plants. |

**Table 2.** Summary of abundance and richness values across the 90 sampling points in forest, cashew and rubber.

|  | Forest | Cashew | Rubber |
| --- | --- | --- | --- |
| Number of points | 30 | 30 | 30 |
| Replicates per point | 6 | 6 | 6 |
| Recorded species richness | 76 | 62 | 58 |
| Total number of individuals | 1028 | 1204 | 605 |
| Mean number of individuals per point (SE) | 5.71 (0.31) | 6.69 (0.46) | 3.36 (0.22) |
| Pooled species richness per point (SE) | 15.1 (0.69) | 14 (0.77) | 9.03 (0.54) |
| Number of species unique to habitat | 22 | 9 | 8 |
| Number of Western Ghat endemics | 10 | 2 | 7 |

**Table 3.**
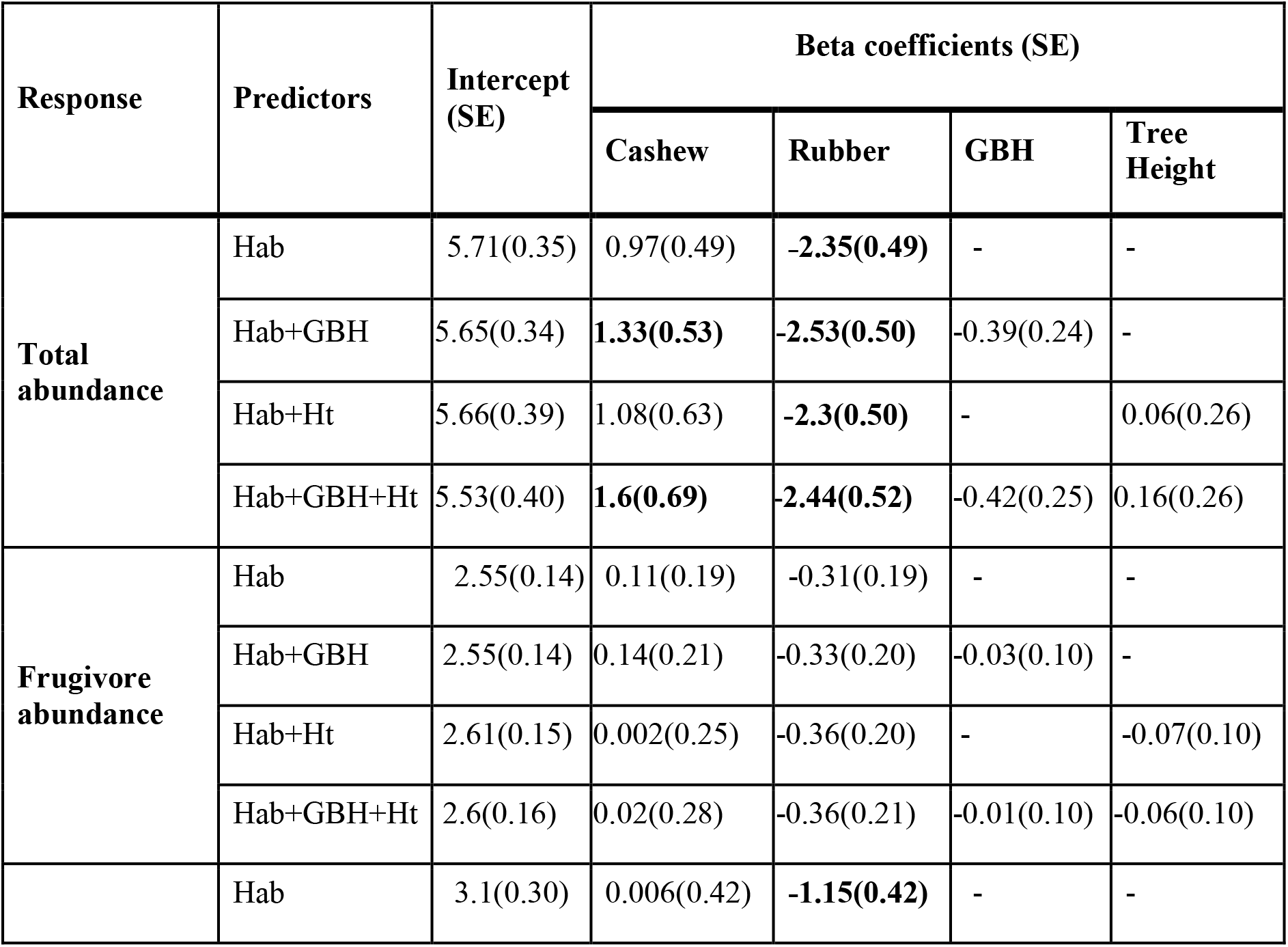
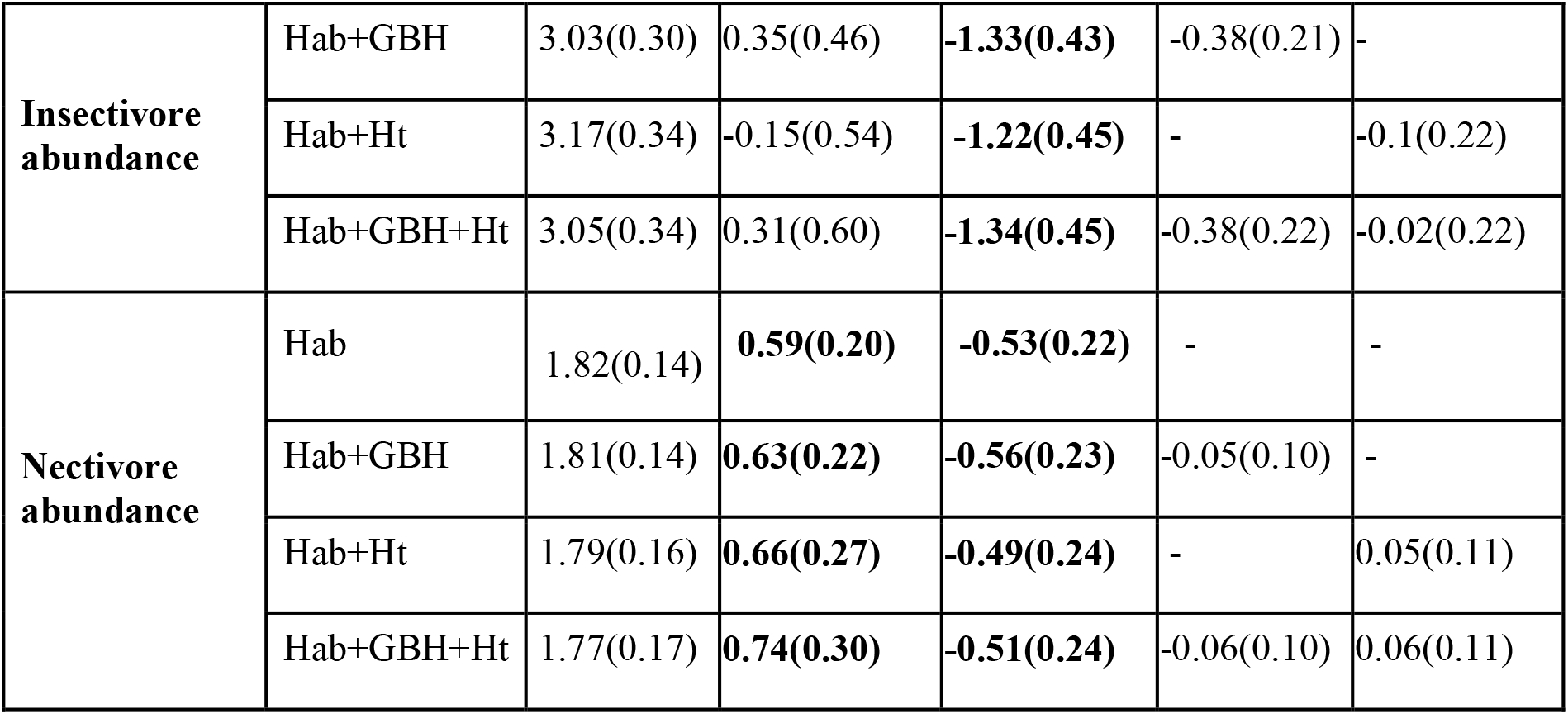
Influence of predictors in four models on total bird abundance and guild-based abundances across 6 sampling sessions. Forest habitat is the intercept. Beta coefficients in which zero does not fall within 95% confidence intervals around the mean are given in bold. Legend: Hab-Habitat; GBH-Tree girth; Ht -Tree height

**Table 4.** Influence of distance to the nearest forest edge on total and guild-based bird abundances in cashew and rubber. Forest habitat is the intercept. Beta coefficients in which zero does not fall within 95% confidence intervals around the mean are given in bold. Legend: Dist-Distance to nearest forest edge

| **Habitat** | **Response** | **Predictor** | **Intercept (SE)** | **Dist (SE)** |
| --- | --- | --- | --- | --- |
| **Cashew** | Total abundance | Dist | 6.67(0.47) | 0.20(0.51) |
|  | Frugivore abundance | Dist | 2.67(0.16) | -0.007(0.17) |
|  | Insectivore abundance | Dist | 3.06(0.26) | **0.72(0.28)** |
|  | Nectivore abundance | Dist | 2.44(0.19) | **-0.45(0.20)** |
| **Rubber** | Total abundance | Dist | 3.38(0.22) | 0.25(0.21) |
|  | Frugivore abundance | Dist | 2.24(0.12) | 0.02(0.11) |
|  | Insectivore abundance | Dist | 1.97(0.39) | 0.22(0.37) |
|  | Nectivore abundance | Dist | 1.29(0.10) | -0.08(0.09) |

#### Richness parameters

Model results (Table 5) revealed substantially lower overall species richness in rubber compared to forest (p<0.001). On the other hand, cashew did not have significantly different species richness compared to forest.

**Table 5.** Influence of predictors on total bird richness and guild-based richness across 6 sampling sessions. Forest habitat is the intercept. Beta coefficients in which zero does not fall within 95% confidence intervals around the mean are given in bold. Legend: Hab-Habitat; GBH-Tree girth; Ht -Tree height

| Response | Model | Intercept (SE) | Beta coefficients (SE) |  |  |  |
| --- | --- | --- | --- | --- | --- | --- |
|  |  |  | Cashew (SE) | Rubber (SE) | GBH (SE) | Tree height (SE) |
| **Total species richness** | Hab | 2.71(0.04) | -0.07(0.06) | **-0.51(0.07)** | - | - |
|  | Hab+GBH | 2.70(0.04) | -0.02(0.07) | **-0.53(0.07)** | -0.05(0.03) | - |
|  | Hab+Ht | 2.69(0.05) | -0.04(0.08) | **-0.50(0.08)** | - | 0.02(0.03) |
|  | Hab+GBH+Ht | 2.68(0.05) | 0.03(0.09) | **-0.51(0.08)** | -0.06(0.03) | 0.03(0.03) |
| **Frugivore species richness** | Hab | 1.64(0.08) | -0.19(0.11) | **-0.38(0.12)** | - | - |
|  | Hab+GBH | 1.63(0.08) | -0.16(0.13) | **-0.4(0.13)** | -0.02(0.06) | - |
|  | Hab+Ht | 1.63(0.09) | -0.18(0.15) | **-0.38(0.13)** | - | 0.008(0.06) |
|  | Hab+GBH+Ht | 1.62(0.09) | -0.14(0.17) | **-0.39(0.13)** | -0.03(0.06) | 0.01(0.06) |
| **Insectivore species richness** | Hab | 2.01(0.06) | -0.16(0.09) | **-0.76(0.12)** | - | - |
|  | Hab+GBH | 1.99(0.06) | -0.08(0.10) | **-0.81(0.12)** | -0.09(0.05) | - |
|  | Hab+Ht | 2.00(0.07) | -0.14(0.12) | **-0.75(0.12)** | - | 0.01(0.05) |
|  | Hab+GBH+Ht | 1.96(0.08) | -0.01(0.14) | **-0.78(0.12)** | -0.10(0.05) | 0.04(0.05) |
| **Nectivore species richness** | Hab | 0.54(0.14) | 0.09(0.19) | -0.13(0.23) | - | - |
|  | Hab+GBH | 0.52(0.14) | 0.16(0.21) | -0.17(0.23) | -0.07(0.10) | - |
|  | Hab+Ht | 0.48(0.16) | 0.21(0.26) | -0.07(0.24) | - | 0.08(0.10) |
|  | Hab+GBH+Ht | 0.45(0.16) | 0.32(0.28) | -0.10(0.24) | -0.09(0.10) | 0.10(0.11) |

For frugivores, all models indicated significantly lower species richness in rubber compared to forest (p<0.01). Similarly, all models showed significantly lower insectivore species richness in rubber compared to forest (p<0.001). Nectivore richness, however, did not show significant difference between any of the three habitat types.

Models indicated no significant effect of tree girth and tree height on the response variables. Distance to the nearest forest edge also had no significant effect on species richness in cashew and rubber (Table 6, Fig. 7.).

**Table 6.** Influence of distance to the nearest forest edge on total and guild-based bird richness in cashew and rubber across 6 sampling sessions. Forest habitat is the intercept. Beta coefficients in which zero does not fall within 95% confidence intervals around the mean are given in bold. Legend: Dist-Distance to nearest forest edge

| Habitat | Response | Model | Intercept (SE) | Dist (SE) |
| --- | --- | --- | --- | --- |
| Cashew | Total species richness | Dist | 2.63(0.04) | 0.04(0.05) |
|  | Frugivore species richness | Dist | 1.45(0.08) | -0.02(0.09) |
|  | Insectivore species richness | Dist | 1.82(0.07) | 0.12(0.07). |
|  | Nectivore species richness | Dist | 0.63(0.47) | 0.20(0.51) |
| **Rubber** | Total species richness | Dist | 2.20(0.06) | 0.02(0.05) |
|  | Frugivore species richness | Dist | 1.25(0.09) | 0.02(0.08) |
|  | Insectivore species richness | Dist | 1.24(0.10) | 0.01(0.09) |
|  | Nectivore species richness | Dist | 0.39(0.18) | -0.09(0.19) |

**Fig. 7.**
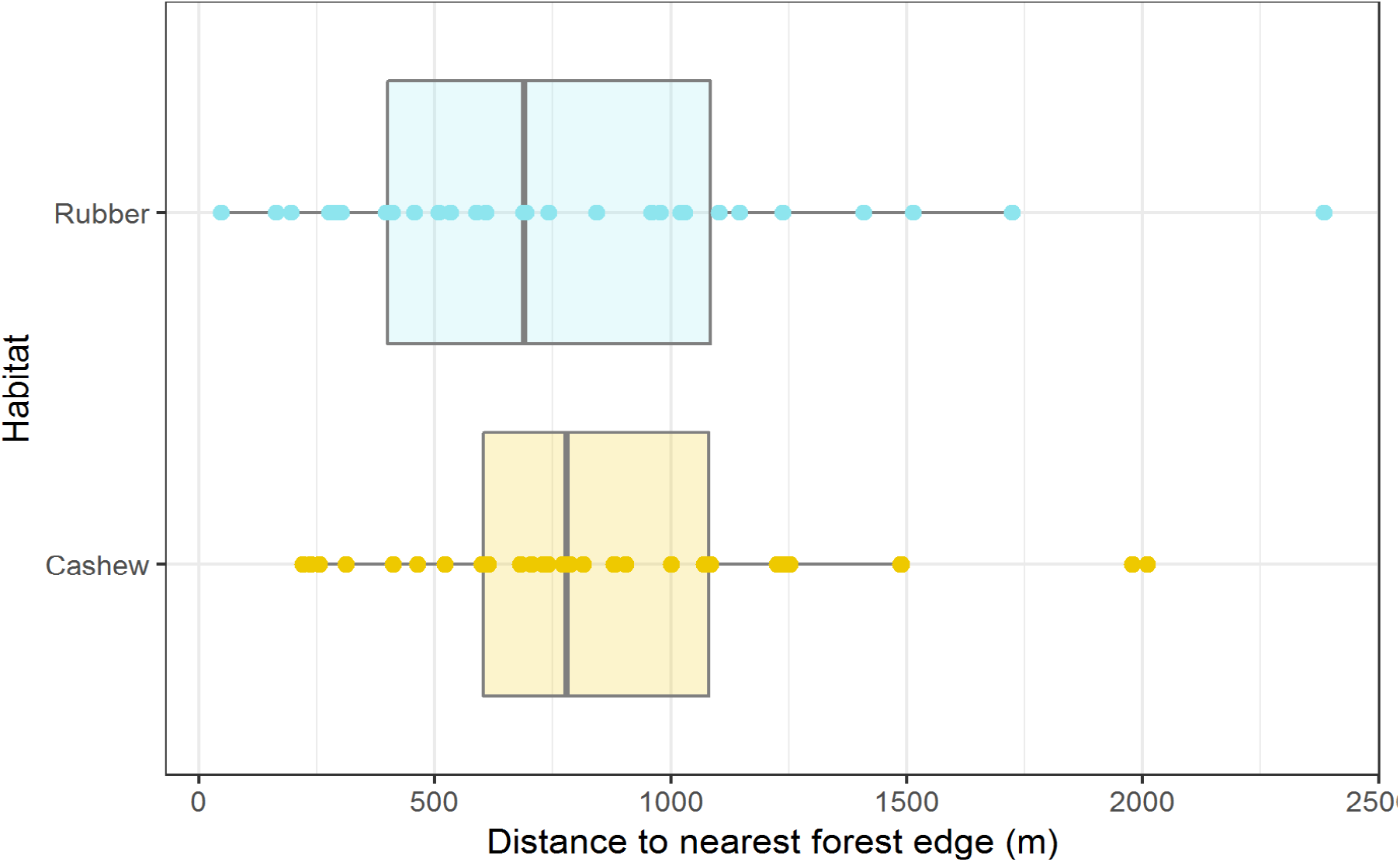
Box and whiskers plot showing distances of all the sampling points in rubber and cashew from the nearest forest edge.

## DISCUSSION

In this study we set out to compare the community structure of birds in cashew and rubber plantations with that in forests in the same landscape, and to examine the influence of habitat attributes on bird communities in cashew and rubber plantations. Our study shows that diversity (richness and abundance) in cashew is comparable to nearby forests, but rubber is less diverse, with fewer individuals and species. When the three most abundant guilds were separately examined (insectivores, frugivores and nectivores), rubber still had fewer birds and bird species, but cashew showed contrasting responses. Cashew had more nectivorous birds and fewer insectivorous bird species than the nearby forest. We observed differences in tree girth and height among forest, rubber and cashew (Fig. S3) but when bird diversity was examined against these habitat characteristics additively with habitat, no patterns were observed; we only recorded a marginal decrease in insectivore species richness with increased tree girth in one case. Distance from the nearest forest edge was unimportant in determining diversity in rubber, but in cashew, fewer nectivorous birds and higher insectivorous birds were seen away from the forest edge. During this study, phenology was not taken into consideration while comparing abundances between different guilds. More insights about the high abundances of nectarivores could be obtained in future studies including phenology sampling.

Lower diversity in rubber can largely be attributed to habitat structure (with little to no under-story and mid-story) and a management regime that mandates the removal of non-*Hevea* shrubs and trees and the application of pesticides (Fox et al., 2014). *Hevea* which has dehiscent fruits, does not rely on birds for seed dispersal and the flowers are also too small to support nectivorous birds. Aratrakorn et al., (2006) also found in South-East Asia stark decline (up to 70%) in bird richness in rubber plantations. A global meta-analysis has found that non-timber plantation crops like rubber lose half the diversity of nearby forests which is highest among all other forms of land-use (Warren-Thomas et al., 2015; Chaudhary et al., 2016). Declines in frugivore and insectivore species richness in rubber are also consistent with data from South-East Asia (Aratrakorn et al., 2006).

Cashew is managed very differently. Plantations are abandoned for 8 months of the year, pesticides are rarely used and understory is lightly cleared before the fruiting season to enable fruit collection. The lack of overall decrease in bird diversity in cashew compared to forest is not surprising, given that ‘complex’ plantations are known to have more diversity than structurally ‘simple’ plantations (Najera and Simonetti, 2010). Cashew even had marginally higher abundance of birds compared to nearby forest, probably because sampling coincided with the fruiting/flowering season of cashew. Biodiversity in cashew is otherwise poorly studied, but one study from the Western Ghats found that together with areca plantations, cashew retained up to 90% of the diversity but differed in composition (Ranganathan et al., 2008). Bird diversity in cashew plantations, while under studied, has been examined for their role in pest control, pollination and other ecosystem services (Shyama, 1997; 1998). While no ‘pests’ like Parakeet species were found during this study, they have been known to forage on the cashew nut in other similar landscapes. Rege (2016) found that out of 11 species of large mammals in the forests, 9 were also found in the adjoining cashew plantations in the same Tillari landscape.

Distance of the sampling points to the nearest forest did not seem to influence community structure, contrary to other studies in the Western Ghats (Anand et al., 2008, 2010) and elsewhere (Zhang et al., 2017). However, distance of point locations was between 500 to 1000m from a forest edge, allowing dispersal to play a strong role in maintaining diversity (Fig.7.). Additionally, most forest patches in the study region were large, and these large fragments are known to double the species richness observed in rubber (Zhang et al., 2017)- more isolated rubber plantations are therefore likely to show declines with distance. An important nectivore (and a facultative insectivore) in the region is the Crimson-backed Sunbird (*Leptocoma minima*). The high abundance of Crimson-backed sunbirds coinciding with the flowering and fruiting season of cashew indicates that sunbirds track resources. However, home-range sizes of sunbirds are known to be small, (< 1km^2^) and it is likely that they cannot track resources for long distances, as seen with decreased nectivore abundance in cashew with distance (Tottrup et al., 2004).

My sampling effort was sufficient, except perhaps for rubber (Fig. 2.). However, our primary goal was to keep effort constant. With identical sampling effort, rubber had half as many individual birds as cashew or forest (Table 2). Many birds go undetected in 30-metre fixed-radius point-count sampling. However, visibility was good in rubber (with no under- and mid-storey) and cashew (with short trees and a sparse dry season canopy) but this is not the case in the nearby moist deciduous and semi-evergreen forests. Actual bird diversity in forests is likely to be even higher than what was observed, and thus encounter rates may provide a conservative estimate. It is also important to note that the forests here are designated as reserve forests or private forests, receiving little or no protection from the state. These forests are therefore at various stages of disturbance, depending on access and resources within. Hence it is very likely that these forests harbour lower bird diversity than better protected forests in the northern Western Ghats landscape.

Studies have shown that rubber is only better than oil palm and open ground in terms of biodiversity. Area under rubber is expected to increase four-fold in South-east Asia (Peh et al., 2006; Aratrakorn et al., 2006; Fox et al., 2014). Rubber also has environmental consequences - higher evapo-transpiration from rubber is correlated with reduced dry season flows and water scarcity (Tan et al., 2011; Giambelluca et al., 2016). Although agro-plantations in general have negative impacts on diversity (as compared to the diversity in forests), it was found that mixed-species plantations with native trees, small stands with long rotation cycles within a connected landscape had the least negative impact (Castano-Villa et al., 2019). Interestingly, despite rubber having no understory and that being cited as potential reason for low diversity, the global review found that such characteristics were not as important as the landscape level features (Castano-Villa et al., 2019; Karanth et al., 2016). When conversion of rubber to more biodiversity-friendly crops is not possible, focus may perhaps be better used encouraging mixed-cropping of rubber interspersed with forest, rather than improving habitat structure *per se*.

Cashew plantations have been very poorly studied for biodiversity, and this study provides some baseline information for bird diversity in cashew. Given that cashew is a long-rotation crop with minimal management for much of the year, the patterns of comparable diversity is expected. Decline of nectivore abundance and of pollination success with distance from forest edge from another study, highlights the potential to maximize biodiversity and ecosystem services benefits by ensuring nearby forests within the landscape (Fretas et al., 2014). While this study found no distance effects because of most point locations being embedded in a nearby forest matrix, reviews from the region point to the strong role of distance to forests at a landscape level in maintaining diversity (Anand et al., 2010).

## ACKNOWLEDGEMENTS

I would like to acknowledge Narayan Desai, who assisted me on field. We would also like to express our sincere gratitude to Shri A.S.Patil (CCF-Territorial) and the Maharashtra Forest Department for the permits. We would like to thank Mr. Kudalkar from the Maharashtra Rubber Board and Sachin Manerikar from Konal Katta for helping us get in touch with plantations owners. We are indebted to Dr. Jayashree Ratnam, Chandni Gurusrikar, Dr. Devcharan Jathanna, Dr. Anindya Sinha and Dr. Hari Sridhar for their guidance during various stages. We are thankful to Girish Punjabi, Anushka Rege, Raman Kulkarni, Malhar Indulkar, Makarand Naik, Pravin Desai, Tanaji Desai, Shashank Dalvi, Ashish Nerlekar, Akshay Surendra, Ishika Ramakrishna, Ramya Roopa, T. S. Rasika, Tarun Menon and Jayu Munje for the help on field, at Bangalore or at home. This project was supported by the Department of Atomic Energy, Government of India and the Sir Dorabji Tata Trust. We would like to express our sincere gratitude to the Tata Institute of Fundamental Research (TIFR-Mumbai) and the National Centre for Biological Sciences, Bangalore, for providing academic and institutional support.

## AUTHOR APPROVAL

All authors have seen and approved the contents of the manuscript and this manuscript has not been accepted or published elsewhere.

