## Supplementary Information for "Bird community structure in a mixed forest-production landscape in the northern Western Ghats, India"

### APPENDICES

A1) Histogram showing the number of aural and visual detections in each habitat type

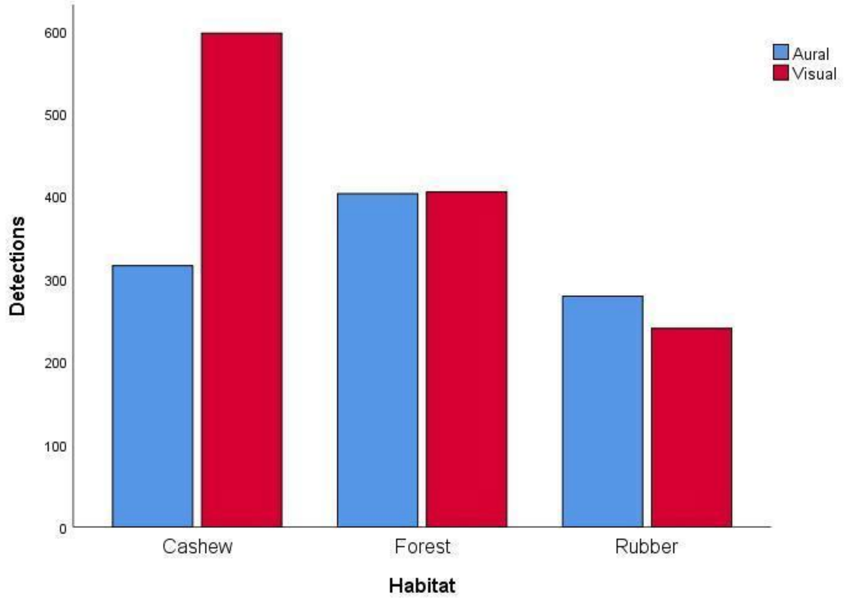


A2) Histograms showing the aural and visual detections in 3 habitats up to 30 metre distance from the point

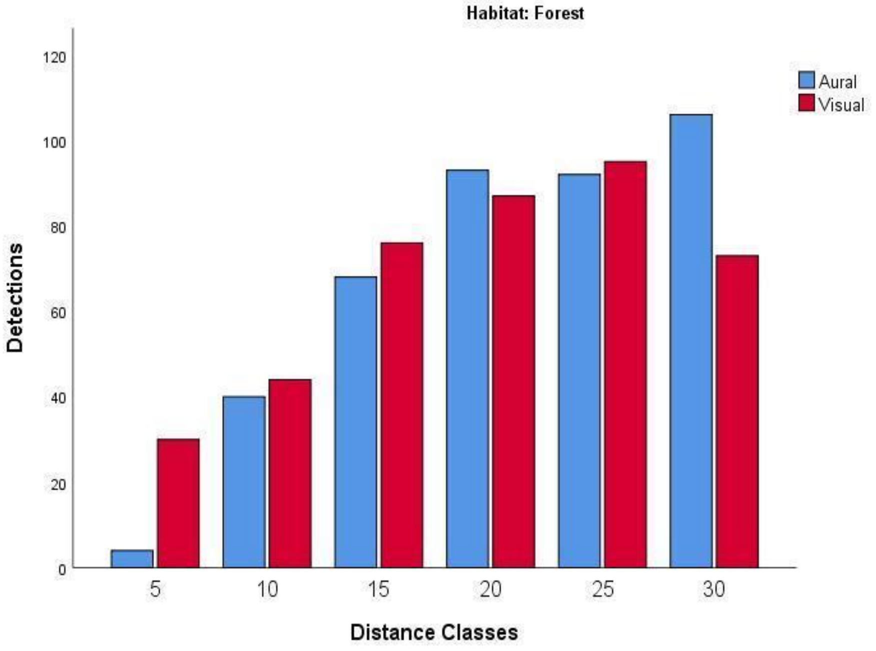

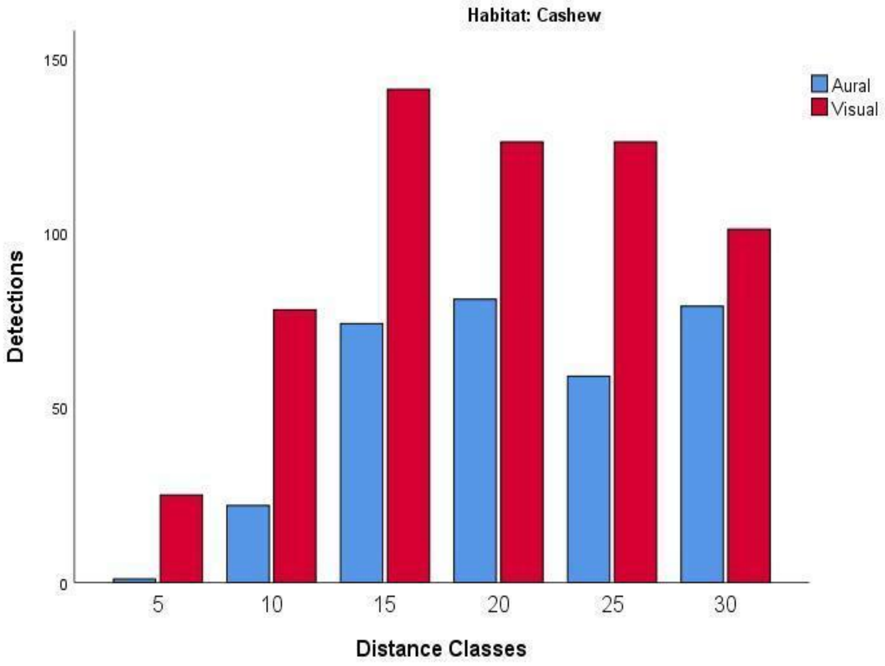
Habitat: Cashew
150
100
50
0
Detections
Aural
Visual
5
10
15
20
25
30
Distance Classes

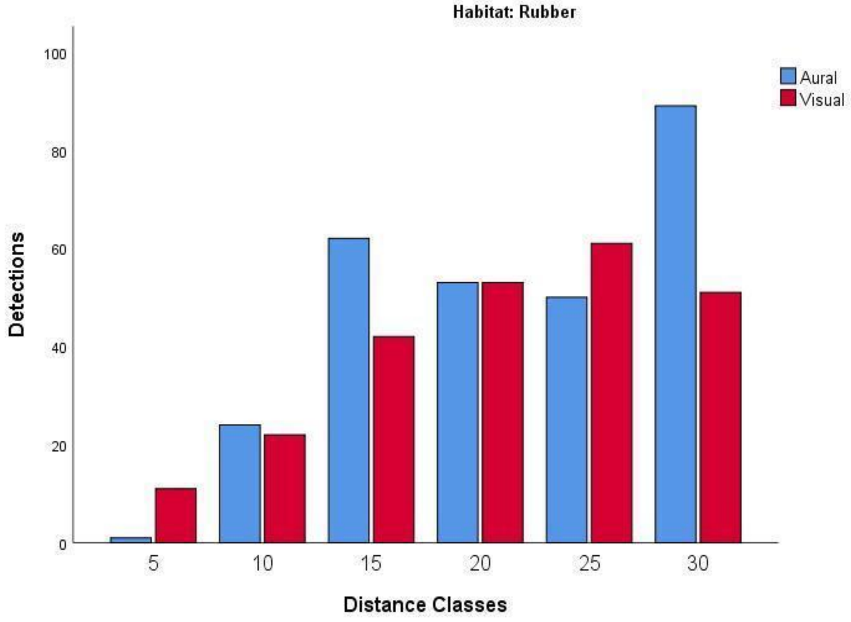
Habitat: Rubber
100
80
60
40
20
0
Detections
Aural
Visual
5
10
15
20
25
30
Distance Classes

A3) Relative abundance of species contributing about 50% of all detected birds in each habitat

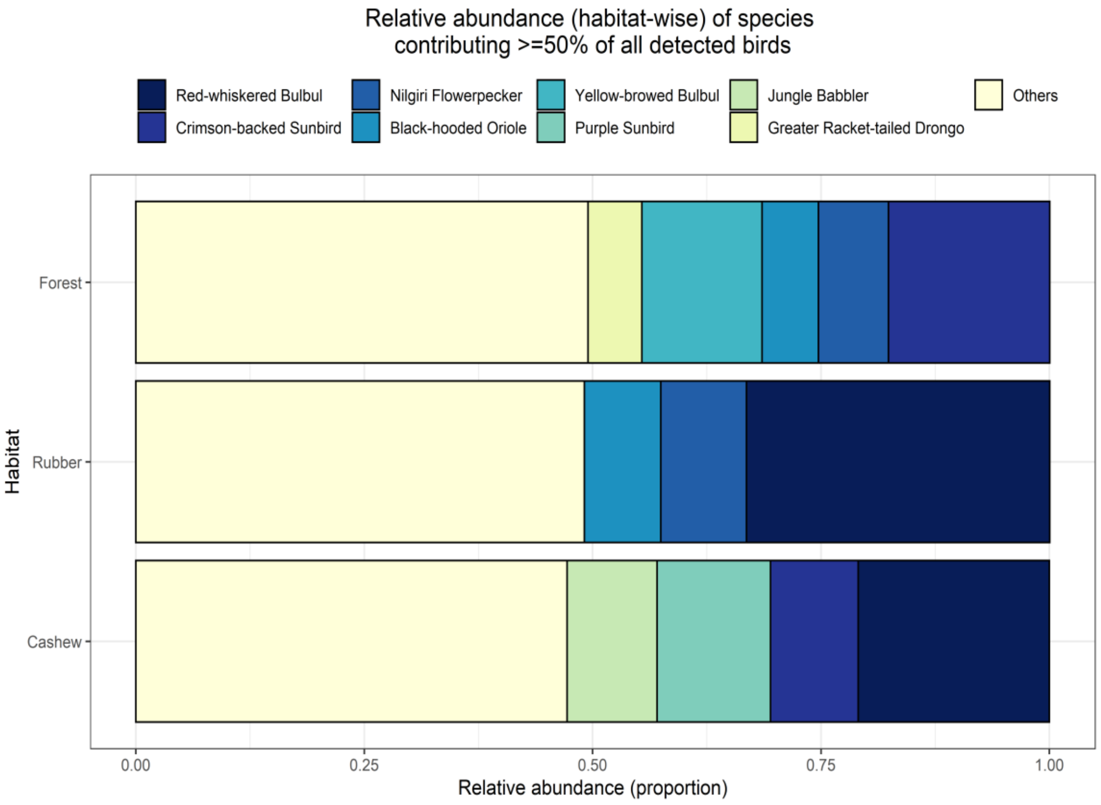


A4) Scatter plot of overall and guild-wise abundances (mean per point) in cashew and rubber with increasing distance to the forest

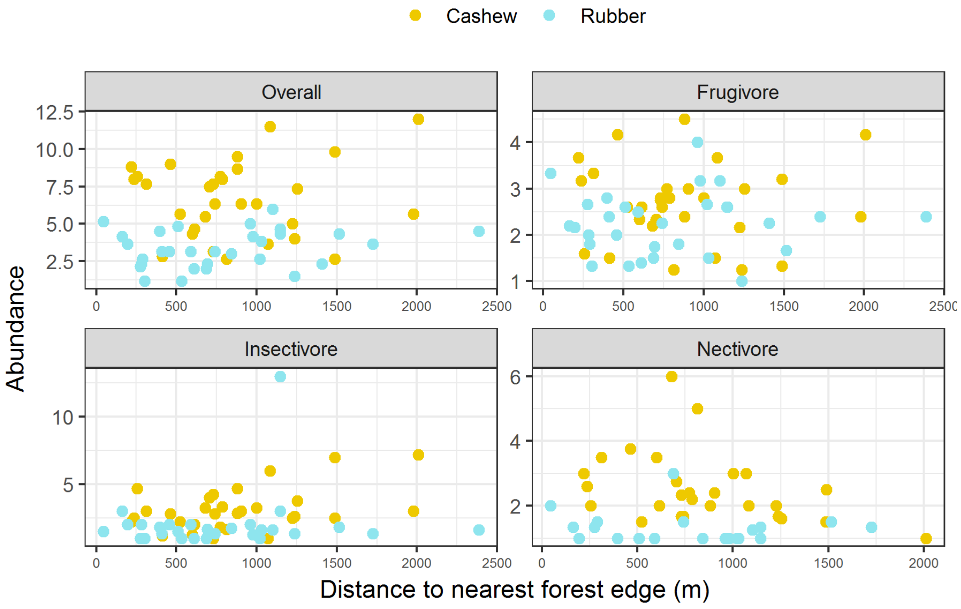

A5) Scatter plot showing overall and guild-wise pooled species richness across points in cashew and rubber with increasing distance to forest

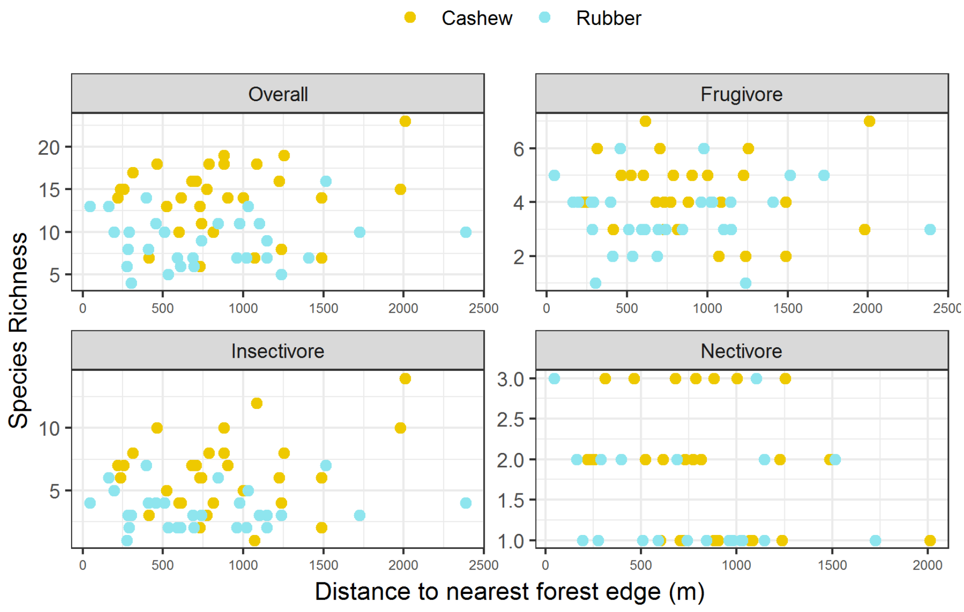

**SUPPLEMENTARY DATA**

S1) Correlogram showing Pearson's correlation coefficient between pairs of predictor variables measured, darker values represent stronger correlations. Legend: m_ht: mean tree height, m_gbh; mean tree girth, m-density: mean tree density, dtnfp: the distance to nearest forest edge, cc: canopy cover and us: understory.

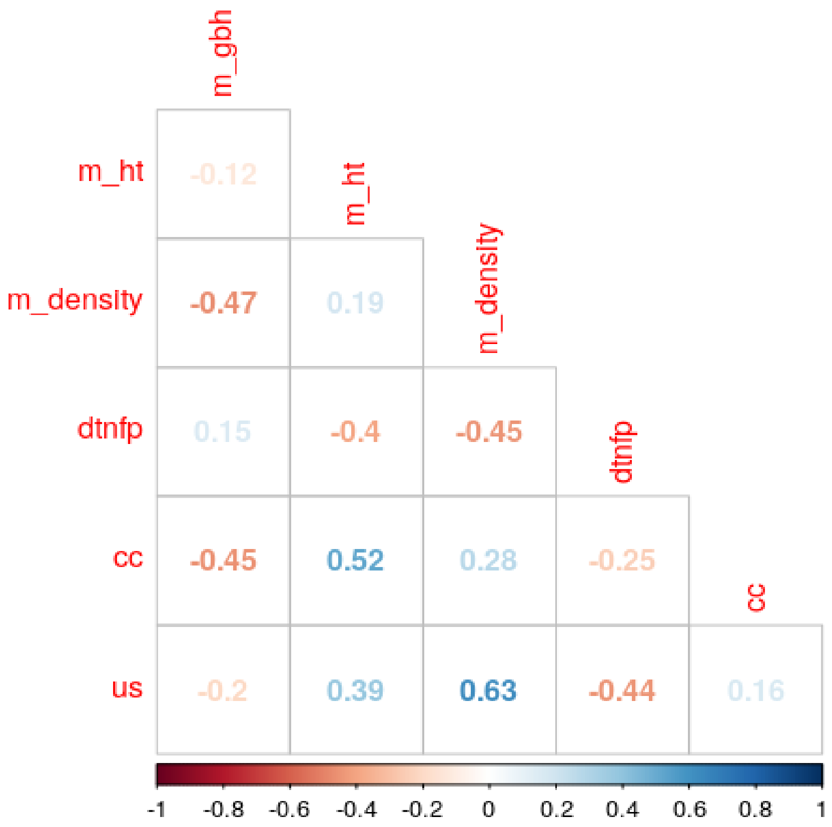

S2) Bird species recorded in point counts during the course of this study along with the dietary guild they belong to, their IUCN status, endemism and the habitats they were observed in (Forest, Cashew and Rubber) (30 locations each and 6 counts per location) (Ali and Ripley, 1978; Rasmussen and Anderton, 2005)

| **Sl No.** | **Common name** | **Scientific Name** | **Family** | **Feeding Guild** | **IUCN Status** | **WG endemic** | **Forest** | **Cashew** | **Rubber** |
|---|---|---|---|---|---|---|---|---|---|
| 1 | Besra | *Accipiter virgatus* | Accipitridae | C | LC | No | Yes | No | No |
| 2 | Crested Serpent Eagle | *Spilornis cheela* | Accipitridae | C | LC | No | Yes | No | No |
| 3 | Shikra | *Accipiter badius* | Accipitridae | C | LC | No | Yes | No | No |
| 4 | Blyth's Reed Warbler | *Acrocephalus dumetorum* | Acrocephalidae | I | LC | No | Yes | Yes | Yes |
| 5 | Common Iora | *Aegithina tiphia* | Aegithinidae | I | LC | No | Yes | Yes | Yes |
| 6 | White-throated Kingfisher | *Halcyon smyrnensis* | Alcedinidae | C | LC | No | No | Yes | Yes |
| 7 | Cattle Egret | *Bubulcus ibis* | Ardeidae | W | LC | No | No | Yes | No |
| 8 | Indian Pond Heron | *Ardeola grayii* | Ardeidae | W | LC | No | No | No | Yes |
| 9 | Great Hornbill | *Buceros bicornis* | Bucerotidae | F | VU | No | Yes | No | No |
| 10 | Malabar Grey Hornbill | *Ocyceros griseus* | Bucerotidae | F | LC | Yes | Yes | No | Yes |
| 11 | Malabar Pied Hornbill | *Anthracoceros coronatus* | Bucerotidae | F | LC | No | Yes | Yes | No |
| 12 | Black-headed Cuckooshrike | *Lalage melanoptera* | Campephagidae | I | LC | No | Yes | No | No |
| 13 | Large Cuckooshrike | *Coracina macei* | Campephagidae | I | LC | No | Yes | Yes | Yes |
| 14 | Scarlet Minivet | *Pericrocotus flammeus* | Campephagidae | I | LC | No | Yes | Yes | Yes |
| 15 | Small Minivet | *Pericrocotus cinnamomeus* | Campephagidae | I | LC | No | Yes | Yes | No |
| 16 | Golden-fronted Leafbird | *Chloropsis aurifrons* | Chloropseidae | F | LC | No | Yes | Yes | Yes |
| 17 | Ashy Prinia | *Prinia socialis* | Cisticolidae | I | LC | No | No | Yes | No |
| 18 | Common Tailorbird | *Orthotomus sutorius* | Cisticolidae | I | LC | No | Yes | Yes | Yes |

| Sl No. | Common name | Scientific Name | Family | Feeding Guild | IUCN Status | WG endemic | Forest | Cashew | Rubber |
|---|---|---|---|---|---|---|---|---|---|
| 19 | Grey-breasted Prinia | *Prinia hodgsonii* | Cisticolidae | I | LC | No | Yes | Yes | Yes |
| 20 | Asian Emerald Dove | *Chalcophaps indica* | Columbidae | G | LC | No | Yes | Yes | Yes |
| 21 | Grey-fronted Green Pigeon | *Treron affinis* | Columbidae | F | LC | Yes | Yes | No | Yes |
| 22 | Mountain Imperial Pigeon | *Ducula badia* | Columbidae | F | LC | No | Yes | No | No |
| 23 | Oriental Turtle Dove | *Streptopelia orientalis* | Columbidae | G | LC | No | Yes | No | No |
| 24 | Rock Pigeon | *Columba livia* | Columbidae | G | LC | No | No | Yes | No |
| 25 | Spotted Dove | *Stigmatopelia chinensis* | Columbidae | G | LC | No | Yes | Yes | Yes |
| 26 | House Crow | *Corvus splendens* | Corvidae | C | LC | No | No | Yes | No |
| 27 | Large-billed Crow | *Corvus macrorhynchos* | Corvidae | C | LC | No | Yes | Yes | Yes |
| 28 | Rufous Treepie | *Dendrocitta vagabunda* | Corvidae | O | LC | No | Yes | Yes | Yes |
| 29 | Asian Koel | *Eudynamys scolopaceus* | Cuculidae | F | LC | No | No | Yes | Yes |
| 30 | Banded Bay Cuckoo | *Cacomantis sonneratii* | Cuculidae | I | LC | No | Yes | No | No |
| 31 | Greater Coucal | *Centropus sinensis* | Cuculidae | I | LC | No | Yes | Yes | Yes |
| 32 | Nilgiri Flowerpecker | *Dicaeum concolor* | Dicaeidae | F | LC | Yes | Yes | Yes | Yes |
| 33 | Thick-billed Flowerpecker | *Dicaeum agile* | Dicaeidae | F | LC | No | Yes | Yes | No |
| 34 | Ashy Drongo | *Dicrurus leucophaeus* | Dicruridae | I | LC | No | Yes | Yes | Yes |
| 35 | Black Drongo | *Dicrurus macrocercus* | Dicruridae | I | LC | No | Yes | Yes | Yes |
| 36 | Bronzed Drongo | *Dicrurus aeneus* | Dicruridae | I | LC | No | Yes | Yes | Yes |
| 37 | Greater Racket-tailed Drongo | *Dicrurus paradiseus* | Dicruridae | I | LC | No | Yes | Yes | Yes |

| Sl No. | Common name | Scientific Name | Family | Feeding Guild | IUCN Status | WG endemic | Forest | Cashew | Rubber |
|---|---|---|---|---|---|---|---|---|---|
| 38 | White-bellied Drongo | *Dicrurus caerulescens* | Dicruridae | I | LC | No | Yes | Yes | No |
| 39 | Common Rosefinch | *Carpodacus erythrinus* | Fringillidae | G | LC | No | No | Yes | No |
| 40 | Asian Fairy-bluebird | *Irena puella* | Irenidae | I | LC | No | Yes | No | Yes |
| 41 | Jungle Babbler | *Turdoides striata* | Leiothrichidae | I | LC | No | No | Yes | Yes |
| 42 | Brown-headed Barbet | *Psilopogon zeylanicus* | Megalaimidae | F | LC | No | Yes | No | No |
| 43 | Coppersmith Barbet | *Psilopogon haemacephalus* | Megalaimidae | F | LC | No | Yes | No | No |
| 44 | Malabar Barbet | *Psilopogon malabarica* | Megalaimidae | F | LC | Yes | Yes | No | No |
| 45 | White-cheeked Barbet | *Megalaima viridis* | Megalaimidae | F | LC | No | Yes | Yes | Yes |
| 46 | Chestnut-headed Bee-eater | *Merops leschenaulti* | Meropidae | I | LC | No | Yes | Yes | Yes |
| 47 | Green Bee-eater | *Merops orientalis* | Meropidae | I | LC | No | No | Yes | Yes |
| 48 | Indian Paradise-flycatcher | *Terpsiphone paradisi* | Monarchidae | I | LC | No | Yes | Yes | Yes |
| 49 | Black-naped Monarch | *Hypothymis azurea* | Monarchidae | I | LC | No | Yes | Yes | Yes |
| 50 | Forest Wagtail | *Dendronanthus indicus* | Motacillidae | I | LC | No | No | No | Yes |
| 51 | Grey Wagtail | *Motacilla cinerea* | Motacillidae | I | LC | No | No | No | Yes |
| 52 | Malabar Whistling Thrush | *Myophonus horsfieldii* | Muscicapidae | I | NT | No | Yes | No | Yes |
| 53 | Oriental Magpie Robin | *Copsychus saularis* | Muscicapidae | I | LC | No | Yes | Yes | Yes |
| 54 | Tickell's Blue Flycatcher | *Cyornis tickelliae* | Muscicapidae | I | LC | No | Yes | Yes | Yes |
| 55 | White-bellied Blue Flycatcher | *Cyornis pallidipes* | Muscicapidae | I | LC | Yes | Yes | No | No |
| 56 | White-rumped Shama | *Kittacincla malabaricus* | Muscicapidae | I | LC | No | Yes | Yes | No |

| Sl No. | Common name | Scientific Name | Family | Feeding Guild | IUCN Status | WG endemic | Forest | Cashew | Rubber |
|---|---|---|---|---|---|---|---|---|---|
| 57 | Vigor's Sunbird | *Aethopyga vigorsii* | Nectariniidae | N | LC | Yes | No | No | Yes |
| 58 | Crimson-backed Sunbird | *Leptocoma minima* | Nectariniidae | N | LC | Yes | Yes | Yes | Yes |
| 59 | Little Spiderhunter | *Arachnothera longirostra* | Nectariniidae | N | LC | No | Yes | Yes | Yes |
| 60 | Purple Sunbird | *Cinnyris asiaticus* | Nectariniidae | N | LC | No | Yes | Yes | Yes |
| 61 | Purple-rumped Sunbird | *Leptocoma zeylonica* | Nectariniidae | N | LC | No | Yes | Yes | Yes |
| 62 | Black-hooded Oriole | *Oriolus chinensis* | Oriolidae | F | LC | No | Yes | Yes | Yes |
| 63 | Indian Golden Oriole | *Oriolus kundoo* | Oriolidae | F | LC | No | Yes | Yes | Yes |
| 64 | Yellow-throated Sparrow | *Gymnoris xanthocollis* | Passeridae | G | LC | No | Yes | No | No |
| 65 | Brown-cheeked Fulvetta | *Alcippe poioicephala* | Pellorneidae | I | LC | No | Yes | Yes | Yes |
| 66 | Puff-throated Babbler | *Pellorneum fuscicapillus* | Pellorneidae | I | LC | No | Yes | Yes | No |
| 67 | Grey Junglefowl | *Gallus sonneratii* | Phasianidae | G | LC | No | No | No | Yes |
| 68 | Indian Peafowl | *Pavo cristatus* | Phasianidae | O | LC | No | No | Yes | No |
| 69 | Red Spurfowl | *Galloperdix spadicea* | Phasianidae | G | LC | No | Yes | No | Yes |
| 70 | Green Leaf Warbler | *Phylloscopus nitidus* | Phylloscopidae | I | LC | No | Yes | Yes | Yes |
| 71 | Greenish Leaf Warbler | *Phylloscopus trochiloides* | Phylloscopidae | I | LC | No | No | No | Yes |
| 72 | Western-crowned Leaf Warbler | *Phylloscopus occipitalis* | Phylloscopidae | I | LC | No | Yes | No | No |
| 73 | Lesser Golden-backed Woodpecker | *Dinopium benghalense* | Picidae | I | LC | No | Yes | Yes | No |

| Sl No. | Common name | Scientific Name | Family | Feeding Guild | IUCN Status | WG endemic | Forest | Cashew | Rubber |
|---|---|---|---|---|---|---|---|---|---|
| 74 | Greater Golden-backed Woodpecker | *Chrysocolaptes guttacristatus* | Picidae | I | LC | No | Yes | No | No |
| 75 | Heart-spotted Woodpecker | *Hemicircus canente* | Picidae | I | LC | No | Yes | Yes | Yes |
| 76 | Rufous Woodpecker | *Micropternus brachyurus* | Picidae | I | LC | No | Yes | No | Yes |
| 77 | Indian Pitta | *Pitta brachyura* | Pittidae | I | LC | No | No | Yes | No |
| 78 | Malabar Parakeet | *Psittacula columboides* | Psittaculidae | F | LC | Yes | No | No | Yes |
| 79 | Plum-headed Parakeet | *Psittacula cyanocephala* | Psittaculidae | F | LC | No | Yes | Yes | No |
| 80 | Vernal Hanging Parrot | *Loriculus vernalis* | Psittaculidae | F | LC | No | Yes | Yes | Yes |
| 81 | Flame-throated Bulbul | *Pycnonotus gularis* | Pycnonotidae | F | LC | Yes | Yes | No | No |
| 82 | Grey-headed Bulbul | *Pycnonotus priocephalus* | Pycnonotidae | F | NT | Yes | Yes | No | Yes |
| 83 | Red-vented Bulbul | *Pycnonotus cafer* | Pycnonotidae | F | LC | No | No | Yes | No |
| 84 | Red-whiskered Bulbul | *Pycnonotus jocosus* | Pycnonotidae | F | LC | No | Yes | Yes | Yes |
| 85 | Yellow-browed Bulbul | *Acritillas indica* | Pycnonotidae | F | LC | No | Yes | Yes | Yes |
| 86 | White-spotted Fantail | *Rhipidura albogularis* | Rhipiduridae | I | LC | No | No | Yes | Yes |
| 87 | Velvet-fronted Nuthatch | *Sitta frontalis* | Sittidae | I | LC | No | Yes | No | No |
| 88 | Brown Wood Owl | *Strix leptogrammica* | Strigidae | C | LC | No | Yes | No | No |
| 89 | Chestnut-tailed Starling | *Sturnia malabarica* | Sturnidae | F | LC | Yes | Yes | No | No |
| 90 | Bar-winged Flycatcher-shrike | *Hemipus picatus* | Tephrodornithidae | I | LC | No | Yes | Yes | No |
| 91 | Common Woodshrike | *Tephrodornis pondicerianus* | Tephrodornithidae | I | LC | No | Yes | Yes | Yes |

| Sl No. | Common name | Scientific Name | Family | Feeding Guild | IUCN Status | WG endemic | Forest | Cashew | Rubber |
|---|---|---|---|---|---|---|---|---|---|
| 92 | Malabar Woodshrike | *Tephrodornis sylvicola* | Tephrodornithidae | I | LC | Yes | Yes | No | No |
| 93 | Dark-fronted Babbler | *Rhopocichla atriceps* | Timaliidae | I | LC | No | Yes | No | No |
| 94 | Indian Scimitar Babbler | *Pomatorhinus horsfieldii* | Timaliidae | I | LC | No | Yes | Yes | No |
| 95 | Tawny-bellied Babbler | *Dumetia hyperythra* | Timaliidae | I | LC | No | No | Yes | No |
| 96 | Malabar Trogon | *Harpactes fasciatus* | Trogonidae | I | LC | No | Yes | No | No |
| 97 | Indian Blackbird | *Turdus simillimus* | Turdidae | I | LC | No | Yes | Yes | Yes |
| 98 | Orange-headed Thrush | *Geokichla citrina* | Turdidae | I | LC | No | Yes | Yes | Yes |
| 99 | Common Barn Owl | *Tylo alba* | Tytonidae | C | LC | No | No | No | Yes |

S3) Scatter plot showing mean tree height and mean tree girth across all the 3 habitats.

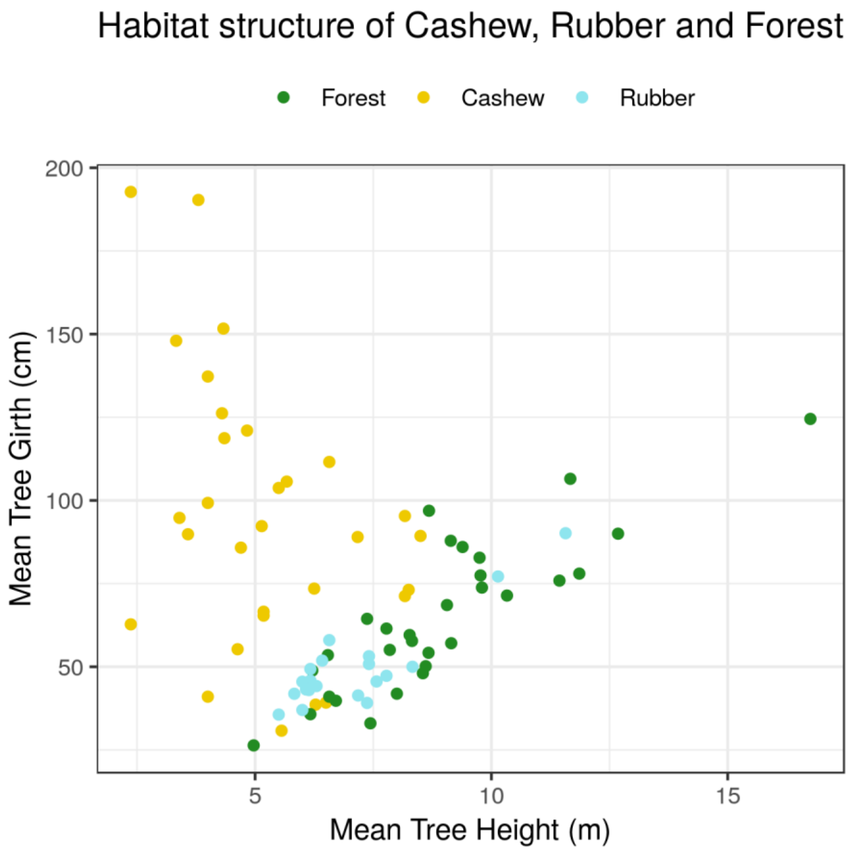
